# Composition and consistence of the bacterial microbiome in upper, middle and lower esophagus before and after Lugol’s iodine staining

**DOI:** 10.1101/375469

**Authors:** Jian Yin, Li Dong, Jing Zhao, Dantong Shao, Anqi liu, Juxiao Li, Aisong Yu, Wenqiang Wei, Wen Chen

**Affiliations:** Department of Epidemiology, National Cancer Center/National Clinical Research Center for Cancer/Cancer Hospital, Chinese Academy of Medical Sciences and Peking Union Medical College, Beijing 100021, China; institutes of Biomedical Sciences, Shanxi University, Taiyuan, 030006, China; School of Public Health, Peking Union Medical College, Beijing 100730, China

**Author notes:** Co-first authors (Jian Yin, Li Dong). Address correspondence to Wen Chen,; Wenqiang Wei.

**Keywords:** Bacteria, Microbiome, 16S rRNA gene, Esophageal microbiome, Lugol’s iodine staining

## Abstract

Esophageal bacteria, as the integral composition of human ecosystem, have been reported to be associated with esophageal lesions. However, few studies focus on microbial compositions in different esophageal segments, especially after Lugol’s iodine staining (LIS) in the endoscopic examination for the screening of esophageal cancer. To investigate the composition of the bacterial microbiome in upper, middle and lower esophagus and if LIS would affect the detection of bacteria, 141 fasting samples including the upper, middle and lower esophagus from 27 participants were collected by brushing the mucosal surface of the esophagus before (Eso) and after (Lug) LIS. Bacterial V3-V4 region of 16S rRNA gene was amplified and sequenced by Illumina’s sequencing platform and analyzed using LEfSe system to identify specific microbiota. The top six abundant bacterial phyla taxa among three locations from both Eso and Lug groups were Proteobacteria, Firmicutes, Bacteroidetes, Actinobacteria, Fusobacteria and TM7. In terms of genera, the bacterium in three locations from two groups was all characterized by a highest relative abundance of *Streptococcus.* Bacteria diversity and the relative abundance between Eso and Lug were comparable (*P* > 0.05). Bacteria diversity was consistent in different esophageal locations for an individual, but it was significantly distinguishing in different subjects (*P* < 0.05). In Conclusion, the bacterial microbiome in healthy esophagus are highly diverse and consistent even among three physiological stenosis at all clades. Lugol’s iodine staining would not change local microenvironment in term of microbial composition. These finding provide an essential baseline for future studies investigating local and systemic bacterial microbiome and esophageal diseases.

## INTRODUCTION

The bacterial microbiome are integral components in all parts of the human digestive tract, from the oral cavity to the anus. Their balance plays crucial roles in the development of mucosal barrier function against the pathogens and immune responses (1). When the balanced bacterial microbiome were disordered with damage of the mucosal barrier, the dysregulated immune response will result in many diseases, such as obesity (2), type 2 diabetes (3), atherosclerosis (4) and cancers (5-7). Therefore, it is essential to test the bacteria of healthy individual to observe significant variations both in pre-clinical conditions and in disease status to understand disease occurrence and progression.

The esophagus consists of the upper, middle and lower segments. It plays the primary role of transferring the food from the oral cavity to the stomach. The environment of the proximal esophageal mucosa is similar to the oral cavity’s one, which the pH value is usually around 7; the environment of middle esophagus is intermediate between the oral-like one and gastric one; the environment of the distal is more like the gastric one because reflux of gastric materials may occur and cause a sudden lowering of pH values (down to 2) (8). In addition, the incidence and the survival time of esophageal squamous cell carcinoma in upper, middle and lower esophagus are different (9) and the tumor location is an independent factor affecting survival time of patients with esophageal cancer. Since the microenvironment of three segments of esophagus exist differences, bacterial microbiome in three locations may be possibly different.

Moreover, studies have demonstrated that screening and diagnosis for esophageal disease with endoscopic Lugol’s iodine staining (LIS) examination could improve early detection of precancerous lesion (such as dysplasia (10) and intraepithelial neoplasia (11)) and esophageal carcinoma (12), but there are few studies compared the bacterial composition before and after LIS.

In this study, using the high-throughput next-generation sequencing (NGS) to sequence V3-V4 region of 16S rRNA gene of bacteria, we compared the composition and consistence of bacterial microbiome in lower, middle and upper of the healthy esophagus before and after the LIS during endoscopy examination in a population-based esophageal cancer screening.

## MATERIALS AND METHODS

### Subjects Recruitment

Residents in Linzhou city aged between 40 to 69 years old with no contraindications for endoscopic examinations (eg, history of reaction to iodine or lidocaine) and who were mentally and physically competent to provide written informed consent and no consumption of any food or beverage at least 6 hours prior to sample collection were enrolled in Linzhou cancer hospital in Jun 2015. Based on the endoscopy-aided biopsy diagnostics by endoscopists and pathologists, 27 esophageal disease-free individuals were included in the final analyses. This project was approved by Institutional Review Board approval of Cancer Hospital, Chinese Academy of Medical Sciences (Number: 15-048/975).

### Screening Procedure and Sample collection

After the informed and signed consent from the participants were obtained, the socio-demographic information and the history of using antibiotics were surveyed by trained interviewers. With general anesthesia, subjects underwent endoscopy examination by local trained endoscopists. Three mucosal samples from three anatomic locations, the upper third, then the middle third and finally the lower third in order along the esophageal tract were obtained with sterile covered brushes respectively. Thereafter, the Lugol’s iodine (1.2%) solution was used to stain the full length of the esophagus, after 2 minutes, the matched samples from the same three locations were collected as previously. Biopsy were taken at the unstained foci indicating the abnormal lesions and confirmed by pathological diagnosis. All the microbiological samples are preserved in PreservCyt solution (Hologic, Bedford, MA, USA) and transported to the lab on dry ice, and stored in -70 °C refrigerator for use.

### Quality control

The prevention and control of contamination from environment and cross-contamination of bacteria from adjacent habitat sites were crucial to accurately determine the site-specific or inter-individual diversity of the microbiome during the sampling and testing. Besides the strict sterile operation, some measures were taken to control and preclude the possibility of contamination as follows. Firstly, a covered esophageal sampling brush in a protective sheath was used so that it was threaded through the endoscope channel, was deployed at the site of sampling, and was then re-sheathed before being retracted through the endoscope. Secondly, samples collection begun from the upper third, followed with middle third with a new brush and ended at lower third with a new brush to avoid the cross contamination along the endoscope channel surface. Once retracted its end of brush rich with bacterial cells were unsheathed and cut with a sterile scissor before immersed into the cytological preservation solutions and sealed immediately. Finally, to evaluate the environment bacterial microbiome during sample collection, three esophageal sampling brushes as the negative control with the same exposure time in the same environment were tested with the collected samples in the same batch of test. The amount of DNA extracted from three negative controls was beyond the detection limitation of Qubit (< 0.01 ng/μL).

### DNA extraction and MiSeq sequencing of 16S rRNA gene amplicons

DNA was extracted by traditional phenol-chloroform method combining enzymatic, chemical and physical extraction methods and was purified by standard methods (13). DNA density and quality were checked using Qubit and agarose gel electrophoresis (AxyPrepTM DNA Gel Extraction Kit, AXYGEN, CA, USA). Extracted DNA was diluted to 2ng/μL and stored at -20°C for downstream use. Universal primers 5’-GTACTCCTACGGGAGGCAGCA-3’ and 5’-GTGGACTACHVGGGTWTCTAAT-3’ with 8nt barcodes were used to amplify the V3-V4 hypervariable regions of 16S rRNA genes for sequencing using Miseq sequencer. The PCR mixture (25 μL) contained 1x PCR buffer, 1.5 mM MgCl_2_, each deoxynucleoside triphosphate at 0.4 μM, each primer at 1.0 μM and 1 U of TransStart Fast Pfu DNA Polymerase (TransStart®, TransGen Biotech, Beijing, China) and 4 ng genomic DNA. The PCR amplification program included initial denaturation at 94 °C for 3 min, followed by 23 cycles of 94 °C for 30 s, 60 °C for 40 s, and 72 °C for 60 s, and a final extension at 72 °C for 10 min. we conducted three PCR for each sample, and combined them together after PCR amplification. PCR products were subjected to electrophoresis using 1.0% agarose gel. The band with a correct size was excised and purified using Gel Extraction Kit (Omega Bio-tek, USA) and quantified with Qubit. All samples were pooled together with equal molar amount from each sample. The sequencing library was prepared using TruSeq DNA kit (Illumina, CA, USA) according to manufacturer’s instruction. The purified library was diluted, denatured, re-diluted, mixed with PhiX (equal to 30% of final DNA amount) as described in the Illumina library preparation protocols, and then applied to an Illumina Miseq system for sequencing with the Reagent Kit v3 600 cycles (Illumina, CA, USA) as described in the manufacturer’s manual.

### Bioinformatics and statistical analysis

All sequences were processed using the Operational taxonomic unit (OTU) picking (QIIME) pipeline V1.9.1 (14). For each sample, OTUs were selected using open reference OTU picking using the Greengenes database version 13.8 (15) with 97% similarity. Samples were rarefied to 5598 reads (lowest number of reads from all samples). Linear discriminant effect size analysis (LEfSe) (16) was performed using the default parameters at any taxonomic level to find biomarkers differentially represented among the sites in esophagus. For the comparative analyses, we calculated the mean and standard deviation for the alpha diversity metrics by sample type. ANOVA and Student’s t test were used to compare the difference by esophageal sampling location (upper, middle, lower esophagus), use of Lugol’s staining (before/after), and between individuals. The threshold on the logarithmic LDA score for discriminative biomarkers was 2.0. Principal coordinate analysis (PCoA) was applied to ordinate similarity matrices. Distance metrics were used to summarize the overall microbiota variability. Different distance metrics reveal distinctive views of the microbiota structure. We used both non-phylogeny-based distance (Bray-Curtis) and phylogeny-based distance (UniFrac) metrics. The original UniFrac distances include two versions: unweighted UniFrac, which uses OTU presence/absence information, and weighted UniFrac, which is based on the relative abundance OTUs. Unweighted UniFrac is most efficient to capture the variability in community membership as well as rare taxonomic lineages, because the probability of these rare taxa being picked up by sequencing is directly related to their abundance. Weighted UniFrac, on the other hand, is most efficient to capture the variability in the abundant lineages because these lineages contribute the most weight in distance calculations. A generalized version of UniFrac distance has been developed to fill the midpoint (17, 18). We used Pearson’s correlation to evaluate the OTU correlation inter-individual and intra-individual (19). All statistical analyses were conducted using R 3.1.1.

## Results

### Demographic characteristics of subjects

The characteristics of healthy participants in high risk area of esophageal squamous cell carcinoma in China were listed in Table S1. The mean age of subjects was 55.6 ±1.7 years old. Only one participant was ever cigarette smokers and no one alcohol drinkers, and none of the subjects had received antibiotics within at least one months before the investigation (Table S1). Finally, 141 fasting samples in different sections, including the upper, middle and lower esophagus, from 27 healthy subjects prior to (Eso) and after (Lug) esophageal LIS remained for data analysis and were showed in figure 1.

**Figure 1.**
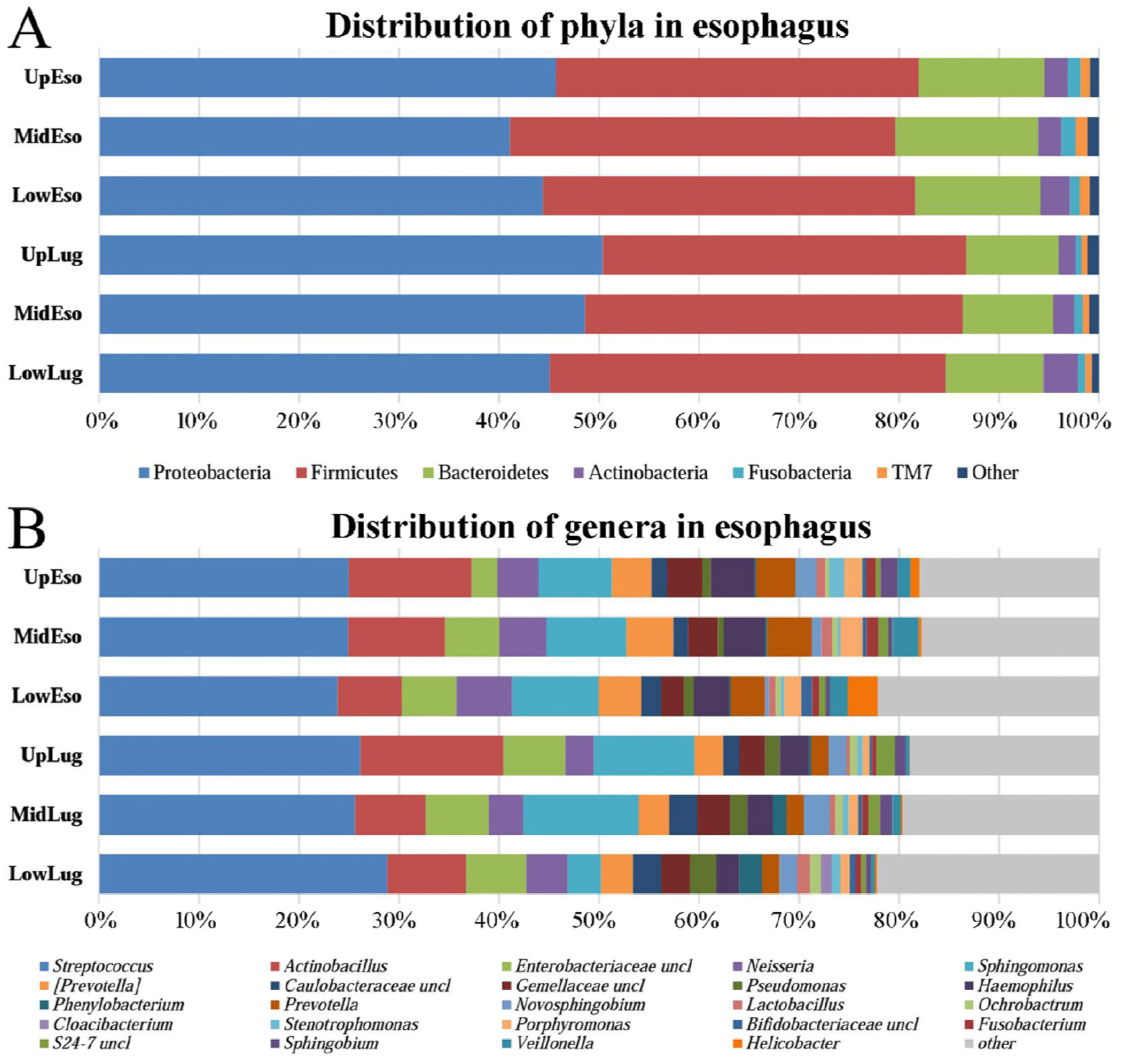
Distribution of Microbiota from three segments of Eso and Lug. Taxonomic composition of the microbiota in the three esophageal habitats investigated based on average relative abundance of 16S rRNA gene next generation sequencing reads assigned to phylum (upper chart: A) and genus (lower chart: B); UpEso, MidEso, LowEso, UpLug, MidLug and LowLug indicate Upper, Middle and Lower esophagus prior to and after Lugol’s iodine staining. Labels indicated genera at average relative abundance > 1 % in at least one body site. The remaining genera were binned together in all phylum as ‘other’ along with the fraction of reads that could not be assigned at the genus level as ‘unclassified’ (uncl). See table S2 for detailed values. Note: Eso indicate the sample from esophagus prior to Lugol’s iodine staining; Lug indicates the sample from esophagus after Lugol’s iodine staining; UpEso indicates upper esophagus; MidEso indicates middle esophagus; LowEso indicates lower esophagus. The addition of Lug at the end of the variable indicates the use of Lugol’s staining for that sample.

### Comparison of microbial communities from esophagus before and after LIS

In the esophagus prior to LIS, the bacterial microbiome at the phyla level in upper esophagus (UpEso) and middle esophagus (MidEso) both consisted of Proteobacteria and Firmicutes followed in decreasing order of relative abundance by Bacteroidetes, Actinobacteria, Fusobacteria and TM7 (Fig. 1a; Table S2); The top four relative abundant bacteria of lower esophagus (LowEso) were same as those of UpEso and MidEso, and the fifth and the sixth relative abundant bacteria were TM7 and Fusobacteria, but the difference between TM7 and Fusobacteria was only 0.10% of relative abundance. In terms of genera, the bacterial microbiome in Eso group was characterized by a highest relative abundance of *Streptococcus,* and *Actinobacillus, Sphingomonas, Neisseria, Haemophilus* and *[Prevotella]* were all over 4% on average in each location of Eso.

In the esophagus after LIS, at the phyla level the top four relative abundant bacterial microbiome of upper esophagus (UpLug) were same as those of UpEso, and the fifth and the sixth bacteria were TM7 and Fusobacteria, but the different value between them was only 0.02%; the top six bacteria in Middle esophagus (MidLug) were identical with those in MidEso; the top four bacteria in Low esophagus (LowLug) were same as those in UpEso, but the fifth and sixth bacteria in LowLug were the reverse order with those in LowEso with different value between them at below 0.10%; Other phyla composed the remaining less than 1% in all locations not only Eso group but also Lug group (Fig. 1a; Table S2). As for genera, the bacterial microbiome in three locations of Lug group was similar to those in three segments of Eso group.

The phyla Proteobacteria, Firmicutes, Bacteroidetes, Actinobacteria were present in at least one esophageal site prior to LIS of all 100% of the subjects, respectively (Table S3). The phyla Fusobacteria and TM7 were present in at least one individual of 100.0%, 93.3% and 100.0%; 93.3%, 100.0% and 100.0% in UpEso, MidEso and LowEso of the subjects. The remaining phyla were all below 87% present in at least one esophageal site prior to LIS and the relative abundance of those were all less than 1% (Tables S2 and S3).

At the genera level, the bacteria with complete prevalence (100%) and high relative abundance (> 1%) were as follows: *Streptococcus, Sphingomonas, Haemophilus, Neisseria, [Prevotella], Prevotella, Veillonella* were present in at least one UpEso, MidEso and LowEso of the subjects. Moreover, the two genera bacteria with high prevalence (> 90%) and high relative abundance (> 1%) were as follows: *Actinobacillus, Porphyromonas* were present in at least one UpEso, MidEso and LowEso of the subjects (Table S3).

Percentages of subjects for which phyla and genera detected in Lug (UpLug, MidLug and LowLug) were nearly same as those in UpEso, MidEso and LowEso, respectively.

### Similar diversity in esophagus microbiome between Eso and Lug, within-Eso and Lug

Calculated microbial diversity indexes of the samples were shown in Table 1. The Chao1, Shannon, and Simpson indexes indicate that species richness and species diversity were not significantly different among samples prior to LIS (all *P* > 0.20), among samples after LIS (all *P* > 0.20), and samples between prior to and after LIS (all *P* > 0.20), after accounting for between individual differences (*P* < 0.05).

**Table 1.** Microbial diversity indices in Eso and Lug

| Samples | Chao1 | Shannon | Simpson |
| --- | --- | --- | --- |
| Eso | 661.9 ± 429.3 | 5.30 ± 1.13 | 0.88 ± 0.10 |
| UpEso | 672.3 ± 362.0 | 5.23 ± 1.06 | 0.88 ± 0.10 |
| MidEso | 645.5 ± 444.1 | 5.33 ± 1.03 | 0.89 ± 0.09 |
| LowEso | 669.9 ± 489.0 | 5.32 ± 1.33 | 0.88 ± 0.12 |
| Lug | 650.2 ± 462.8 | 5.10 ± 1.36 | 0.85 ± 0.14 |
| UpLug | 664.0 ± 488.8 | 4.82 ± 1.31 | 0.84 ± 0.13 |
| MidLug | 671.0 ± 502.7 | 5.24 ± 1.21 | 0.87 ± 0.12 |
| LowLug | 618.0 ± 415.2 | 5.25 ± 1.54 | 0.86 ± 0.17 |

Note: Eso indicates the sample from esophagus prior to Lugol's iodine staining; Lug indicates the sample from esophagus after Lugol's iodine staining; UpEso indicates upper esophagus; MidEso indicates middle esophagus; LowEso indicates lower esophagus. The addition of Lug at the end of the variable indicates the use of Lugol's staining for that sample.

The LEfSe system was used to determine statistically significant biomarkers of clade abundance all taxonomic levels between these groups within the upper digestive tract. When we compared with the samples from three locations in the esophagus prior to LIS, there were not significantly different relative abundance clades. Furthermore, analyzing the influence of LIS for bacterial microbiome, we did not find the significantly different relative abundance clades in all between-group (UpEso and UpLug, MidEso and MidLug, and LowEso and LowLug), and the different abundance also did not present in three locations in esophagus after LIS.

We analyzed the similarity (or diversity) by Bray-Curtis (Fig. 2A), Unweighted UniFrac UniFrac (Fig. 2C) and Weighted UniFrac distance (Fig. 2E), and all the PCoA plots showed separate, large clusters of the esophageal samples; the matched Eso-Lug and the samples of three locations Eso and Lug from an individual were all similar, and the similarity of intraindividual was significantly higher than interindividual based on the Bray-Curtis distance (Fig. 2B), the Unweighted UniFrac (Fig. 2D) and the Weighted UniFrac distance (Fig. 2F), (Fig. 2, all *P* < 1e-14).

**Figure 2.**
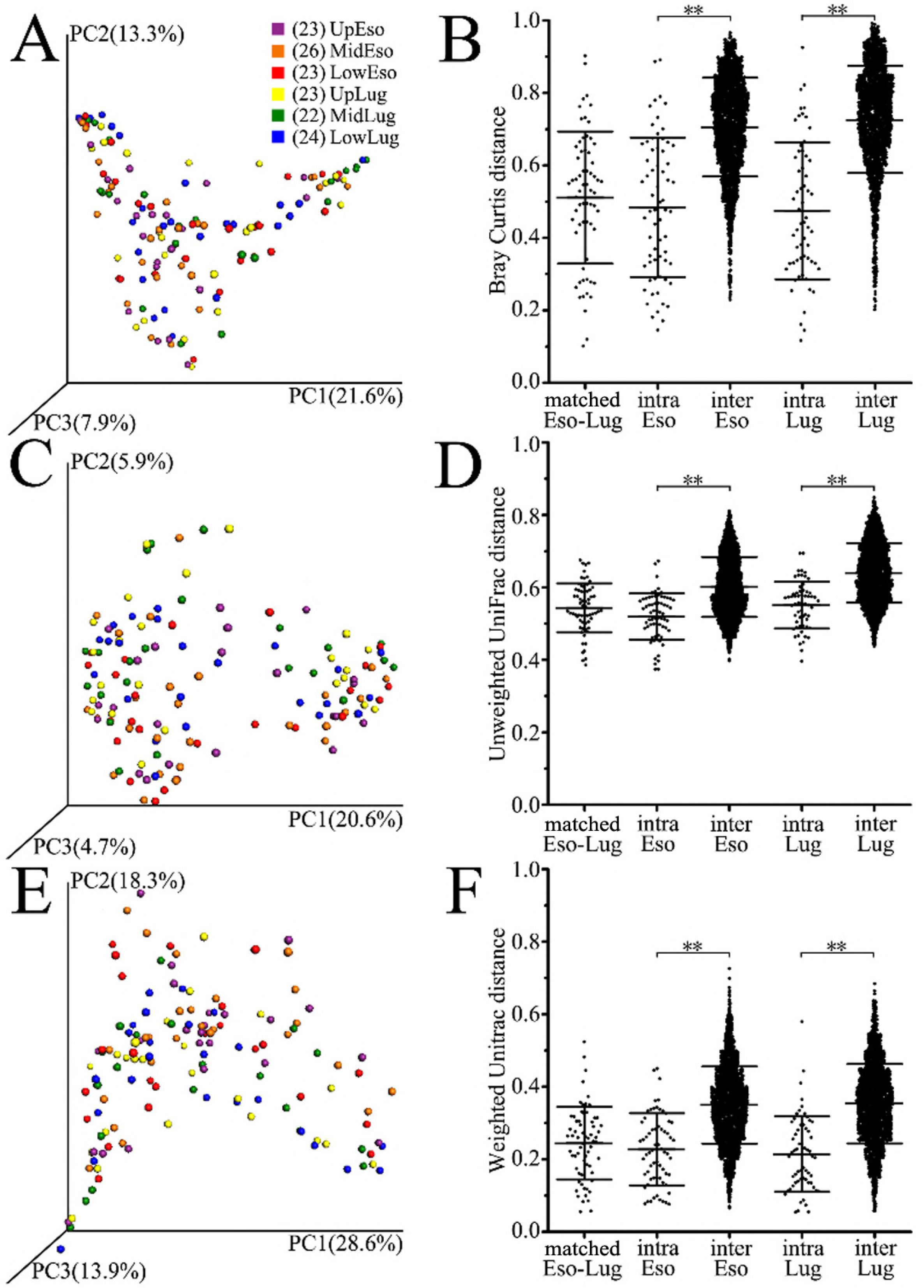
Community structure similarity of intraindividual and interindividual samples in Eso and Lug.
Note: Eso indicates the sample from esophagus prior to Lugol’s iodine staining; Lug indicates the sample from esophagus after Lugol’s iodine staining; Matched Eso-lug indicated the match esophageal samples from prior to and after lugol’s iodine staining; intra Eso indicated the three esophageal samples of individual Eso; inter Eso indicated the samples from different individual Eso; intra Lug indicated the three esophageal samples of individual Lug; inter Lug indicated the samples from different individual Lug.

We further used Pearson’s correlation to evaluate the OTU correlation inter-individual and intra-individual showed in figure 3. We compared the observed OTUs > 1%o of samples from the individuals. The median of significant pearson’s correlations of matched Eso-Lug, intra Eso, inter Eso, intra Lug and inter Lug were 0.82, 0.79, 0.61, 0.93 and 0.59 respectively. The proportions of significant pearson’s correlation great than or equal to 0.99 of matched Eso-Lug, intra Eso and intra Lug were 22.4%, 24.2% and 14.8%, and of the inter Eso and inter Lug were only 1.3% and 3.9%. The proportions of significant pearson’s correlation great than or equal to 0.50 of matched Eso-Lug, intra Eso and intra Lug were 81.0%, 71.0% and 87.0%, and of the inter Eso and inter Lug were only 62.0% and 60.1%. The matched Eso-Lug and three esophageal locations in Eso and Lug from an individual person were all similar, and the similarity of intra-individual was significantly higher than that of inter-individual.

**Figure 3.**
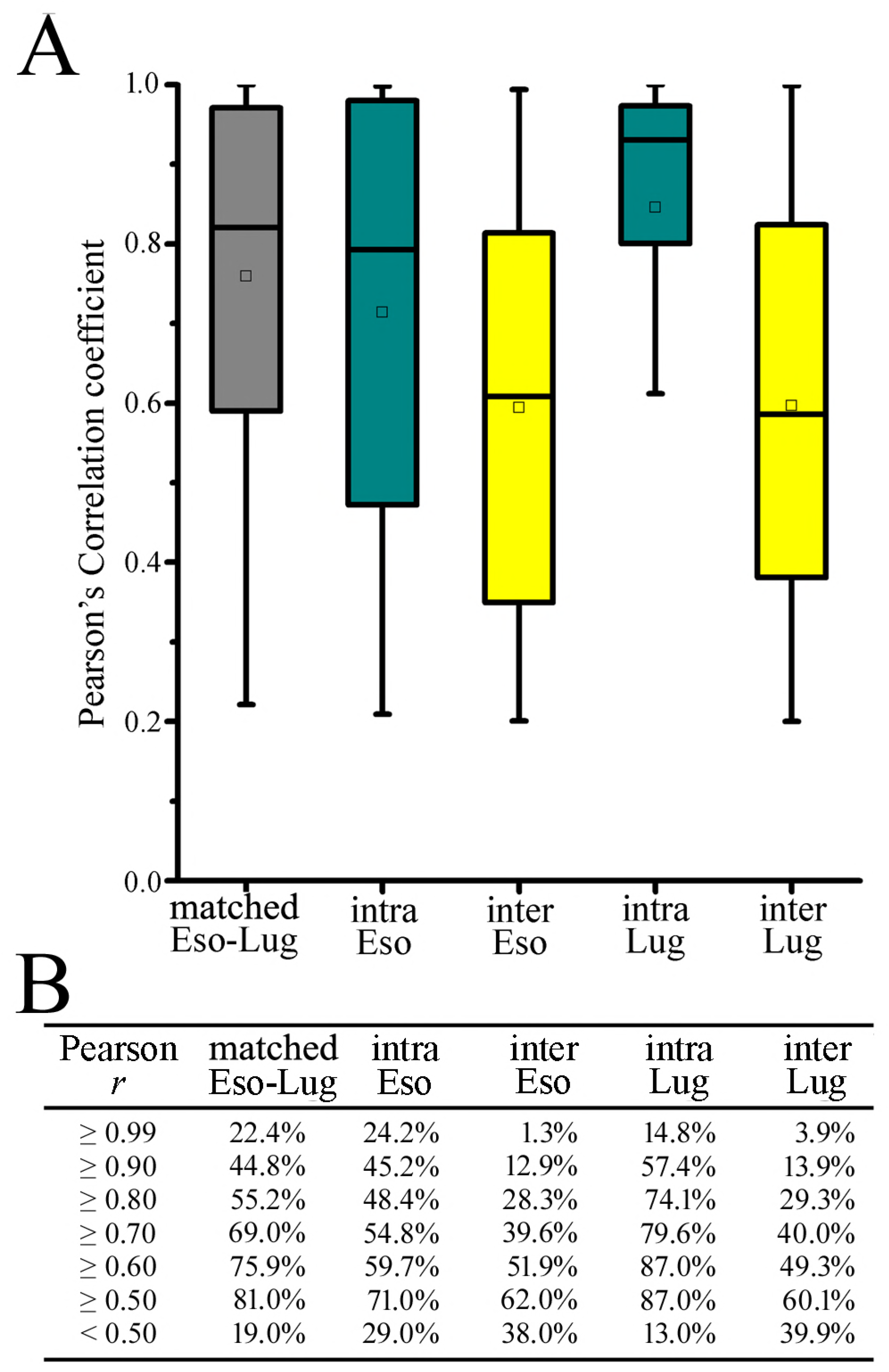
OTU correlation samples between Eso and Lug Note: Eso indicates the sample from esophagus prior to Lugol’s iodine staining; Lug indicates the sample from esophagus after Lugol’s iodine staining; Matched Eso-lug indicated the matched esophageal samples from prior to and after lugol’s iodine staining; intra Eso indicated the three esophageal samples of individual Eso; inter Eso indicated the samples from different individual Eso; intra Lug indicated the three esophageal samples of individual Lug; inter Lug indicated the samples from different individual Lug.

## Discussion

Esophageal cancer is a major upper gastrointestinal malignancy in China and the mortality in China accounts for nearly half of those worldwide according to the report of GLOBACAN in 2012 (20). However, it remains unclear about the biological etiology of esophagus cancer. Since the 1950s, the studies concerning biological causes of esophageal cancer mainly focused on pathogenic fungus (21) and virus (22), particularly single microorganism or several ones (23). Recently, a growing number of studies demonstrated microbial communities play an important role in human physiology and many diseases, in particular those of the digest tract associated with changes in composition and diversity of microbial communities (24). However, basic composition of microbiome in the esophagus is not clear until now. Traditional culture-based methods capture only a small proportion, typically less than 30%, of our bacterial microbiota (7). Culture-independent analysis using next-generation sequencing (NGS) which relies on the amplification and sequencing of the generally considered universal 16S rRNA gene has made up this gap, has been essential in defining and understanding the bacterial microbiome, and greatly has increased appreciation for the complexity hidden in even seemingly simple microbial consortia (25, 26). Even though it has been applying to test bacterial microbiome in human upper gastrointestinal tract, rare is used for detection of bacterial microbiome in the esophagus.

The study as a pilot research clarified firstly and successfully the baseline composition of bacterial microbiome in three esophageal segments, before and after Lugol’s iodine staining using 16S rRNA gene sequencing in healthy population of Linzhou city, a high-risk area of an esophageal cancer, in Henan province of China.

### Bacterial microbiome in upper, middle and lower esophagus

In this study, we identified the relative abundance and presentation of bacterial microbiota of esophageal mucosa samples in three anatomic locations from 27 individuals and found that the consistent distributions of bacterial microbiota of three locations were not only present at the phylum level but genus levels. The most common phyla bacteria were Proteobacteria, Firmicutes, Bacteroidetes, Actinobacteria, Fusobacteria and TM7 in three locations of esophagus, though the order present different, the difference value was very small. Moreover, the six phyla bacteria nearly present all three locations of individual esophagus. Therefore, the distribution of phylum bacteria in three locations of esophagus is similar. Furthermore, the most common genus was *Streptococcus*, followed by the *Actinobacillus*, *Sphingomonas*, *Neisseria*, *Haemophilus and [Prevotella]* in three locations of esophagus in Eso. In addition, there present some different relative abundance of bacteria in three locations in Eso group. These findings on microbiome in esophageal mucosa were basically consistent with those by Pei et al who performed biopsies of distal esophagus from only four patients without esophageal lesions by 16s rRNA gene sequencing (27).

We observed prevalent (present > 90% in at least one in UpEso, MidEso and LowEso) and high abundance of phyla (> 1%) and genera (> 1%) in three locations of esophageal samples prior to LIS. Moreover, the twenty-six phyla whose absolute value of different value of present among UpEso, MidEso and LowEso more than 20%, and the most different value was 46.7%. Therefore, the distribution of genus bacteria in three locations of esophagus is basically consistent.

Given that the different value of bacteria among UpEso, MidEso and LowEso, we further analyzed statistically significant biomarkers of clade abundance all taxonomic levels using LEfSe between different groups within esophageal sites. The biomarkers were not found among three locations of esophagus prior to LIS; Furthermore, richness and diversity were not significantly different among three locations of Eso, but significantly among different subjects (*P* < 0.05).

Based on the findings above, we further compared the similarity (or diversity) in bacteria between intra-individual and inter-individual using the Bray-Curtis, Unweighted Unifrac and Weighted Unifrac measure of beta diversity. Intra-individual distance was very significantly lower (greater similarity) than inter-individual distance (lower similarity) for Eso. We further used Pearson’s correlation to evaluate the OTU correlation inter-individual and intra-individual, the results were same as the Weighted UniFrac distance analysis. Therefore, the bacteria from three locations of esophagus was similar in an individual and was distinguishing from inter-individual; the three locations of esophagus were regarded as an integral whole environment to habitat for bacteria in healthy people.

### The effect of LIS for testing bacteria

Plentiful studies showed that LIS chromoendoscopy is an effective way to boost the detection of esophageal diseases (28), especially precancerous lesion and cancer (29). Moreover, the Lugol’s iodine (1.2%) solution is often used to a medication and disinfectant for numerous purposes. But it is unknown whether the solution affect the identification of bacteria with 16s rRNA gene sequencing. Therefore, we further analyze the microbiome of esophagus after LIS compared with prior to LIS.

The top six phyla bacteria of Lug were same as the bacteria of Eso not only as a whole (Lug vs Eso) but also as tripartite (UpLug vs UpEso; MidLug vs MidEso; LowLug vs LowEso). Moreover, the most common genus microbiome of UpLug, MidLug and LowLug were basically similar to those present in UpEso, MidEso, LowEso. Finally, percentages of subjects for which phyla and genera detected in UpLug, MidLug and LowLug were similar to UpEso, MidEso and LowEso, respectively, except for some different genera bacteria.

Using the LEfSe analysis to detect whether the LIS significantly affects some bacteria relative abundance, we did not find the influenced biomarkers between UpLug and UpEso, MidLug and MidLug, and LowLug and LowEso, and among three locations of esophagus after LIS. Furthermore, richness and diversity were not significantly different among tripartite (UpLug vs UpEso; MidLug vs MidEso; LowLug vs LowEso), and among three locations of Lug, but significantly in different subjects (*P* < 0.05). The results were same as Eso. The results after comparing the beta diversity in every sample measured by the Bray-Curtis, Unweighted Unifrac and Weighted Unifrac showed that the match Eso-Lug and the samples of three locations Lug from the same individual person were all similar, and the similarity of intra-individual were significantly higher than inter-individual for Lug. We further used Pearson’s correlation to evaluate the OTU correlation of matched Eso-Lug, inter-individual and intra-individual, the matched Eso-Lug and three esophageal locations in Eso and Lug from an individual person were all similar, and the similarity of intra-individual were significantly higher than inter-individual. Therefore, the LIS do not significantly affect the detection of microbiome in the esophagus using the high-throughput 16s rRNA gene NGS technologies.

The study has several strengths in stringent inclusion criteria: the subjects were confirmed by physicians aided with esophageal endoscopy to avoid the bias of disease misclassification; a series of quality control methods were taken to minimize of the contamination of microbiota from handling environment and adjacent tracts. There are several limitations of the study needing to be addressed in the future research. Firstly, compared to whole genome shotgun sequencing, microbiome diversity at the species level in high phylogenetic resolution couldn’t be reached by 16S rRNA gene sequencing. Secondly, further large-scale studies are required for validating our findings, especially linking the demographic and clinical characteristics of individuals with the microbial compositions. Finally, since most of participants in our study were women, which might produce microbial bias in term of sex-relevance.

In conclusion, we showed that the bacterial microbiome in normal esophageal was highly diverse and consistent in different sections of esophagus in an individual. The most of high relative abundance bacteria were predominant in the esophagus mucosa. LIS did not significantly affect the bacterial diversity and relative abundance. These data comprehensively provide a critical baseline for future studies investigating the role of microbiome in the local and systemic esophageal diseases affecting human health. Further studies are needed to expand the sample size to validate these findings.

## Acknowledgements

This study was supported by CAMS Innovation Fund for Medical Sciences (CIFMS) (Grant No. 2016-I2M-3-001); Natural Science Fund from the National Natural Science Foundation of China (81573224); Fundamental Research Funds for the Central Universities (2016ZX310178).

